# Unsupervised identification of low-frequency antigen-specific TCRs using distance-based anomaly scoring

**DOI:** 10.64898/2026.03.09.709174

**Authors:** Kyohei Kinoshita, Tetsuya J. Kobayashi

## Abstract

Identifying antigen-specific T cell receptors (TCRs) within the diverse human repertoire remains challenging due to their extremely low frequencies, often as rare as one per million cells. Here, we propose a novel unsupervised approach that detects low-frequency antigen-specific TCRs through distance-based anomaly detection in TCR sequence space. Our method is based on the observation that antigen-specific TCRs preferentially localize at the periphery of V gene clusters rather than cluster centers. Using TCRdist3 to quantify sequence distances, we identify query TCRs that are anomalous compared to reference repertoires within their V-J gene combinations. We validated this approach across three immunological contexts: COVID-19 infection, influenza vaccination, and yellow fever vaccination. For SARS-CoV-2-specific TCR detection in a COVID-19 patient, our method demonstrated 34.3% accuracy, significantly outperforming similarity-based (ALICE: 8.0%) and frequency-based methods (edgeR: 5.8%, the Pogorelyy method: 6.3%), and uniquely detected low-frequency antigen-specific TCRs at clone count one. The minimal overlap with conventional approaches (≤6.7%) indicates our method captures distinct TCR clones overlooked by existing analyses. This spatial distribution-based paradigm provides a complementary strategy for TCR specificity detection, particularly valuable for identifying rare antigen-specific clones essential for understanding immune responses.

## Introduction

T cells play a central role in adaptive immunity by recognizing pathogen-derived peptides via their T cell receptors (TCRs). The human body contains approximately 10^11^ T cells [1], each expressing unique TCRs generated through somatic recombination of variable (V), diversity (D), and joining (J) gene segments [2]. This recombination mechanism, combined with nucleotide deletions and insertions in the complementarity-determining region 3 (CDR3), creates an extraordinarily diverse TCR repertoire capable of recognizing foreign antigens.

Identifying antigen-specific TCRs within this vast repertoire is crucial for understanding immune responses, with direct applications in infectious disease diagnosis, treatment monitoring, and cancer immunotherapy [3–6]. However, the power distribution of TCR frequencies and the extremely low frequency of antigen-specific TCRs, as low as one per million cells, pose a significant challenge for their detection and characterization [7–11].

Current computational approaches to identifying antigen-specific TCRs can be broadly categorized into supervised and unsupervised methods. Supervised deep learning models, including DeepTCR [12], NetTCR [13], and TCR-BERT [14], have demonstrated reliable predictive performance for seen epitopes (those well-represented in the training data).

However, these approaches require large-scale training datasets of TCR-pMHC binding pairs to achieve reasonable prediction accuracy and demonstrate limited generalization to unseen epitopes (those absent from or scarce in the training data), restricting their applicability to novel antigens [15]. This limitation has driven the development of unsupervised approaches independent of TCR-pMHC binding data.

Among unsupervised methods, sequence similarity-based approaches leverage the observation that TCRs with similar sequences often share antigen specificity [16,17]. Tools such as TCRdist3 [18], GLIPH2 [19], and TCRMatch [20] identify groups of functionally related TCRs through sequence clustering or motif detection. However, these methods primarily focus on detecting TCR groups rather than predicting individual TCR specificity. Alternative similarity-based methods, including ALICE [21] and TCRNET [22], detect enrichment of similar TCRs compared to the controls generated from a V(D)J recombination model or healthy donors. However, sequence similarity does not necessarily correspond to functional similarity and often exhibits non-linear relationships, potentially misidentifying TCR specificity [23].

Frequency-based methods represent another class of unsupervised approaches that detect antigen-specific TCRs through statistical analysis of clonal expansion following antigen stimulation. These include adaptations of tools originally developed for differential expression analysis such as edgeR [24] and TCR-specific methods such as the Bayesian framework of Pogorelyy et al. [25]. While effective for highly proliferating clones, these methods cannot detect TCRs that remain at low frequencies due to sequencing errors or limited proliferation capacity. They are particularly limited in contexts with weak immune responses [26], where clonal expansion is minimal. Additionally, these statistical approaches require multiple biological replicates for statistical power, imposing practical constraints on their application.

Here, we introduce TCR-RADAR (Rare Antigen-specific Detection by Anomaly Ranking), a novel unsupervised approach for detecting low-frequency antigen-specific TCRs with high accuracy. Our method is based on the key observation that antigen-specific TCRs exhibit characteristic distribution patterns in sequence space, specifically localizing at the peripheries of V gene clusters. By quantifying the distance between each query TCR and reference TCRs in TCRdist3-defined sequence space, we identify antigen-specific TCRs as anomalies within either their V-J gene combinations or their respective V gene families. TCR-RADAR is fundamentally distinct from existing proliferation-based and similarity-based methods in that it directly employs the spatial distribution characteristics of antigen-specific TCRs **(Fig. 1)**.

**Figure 1.**
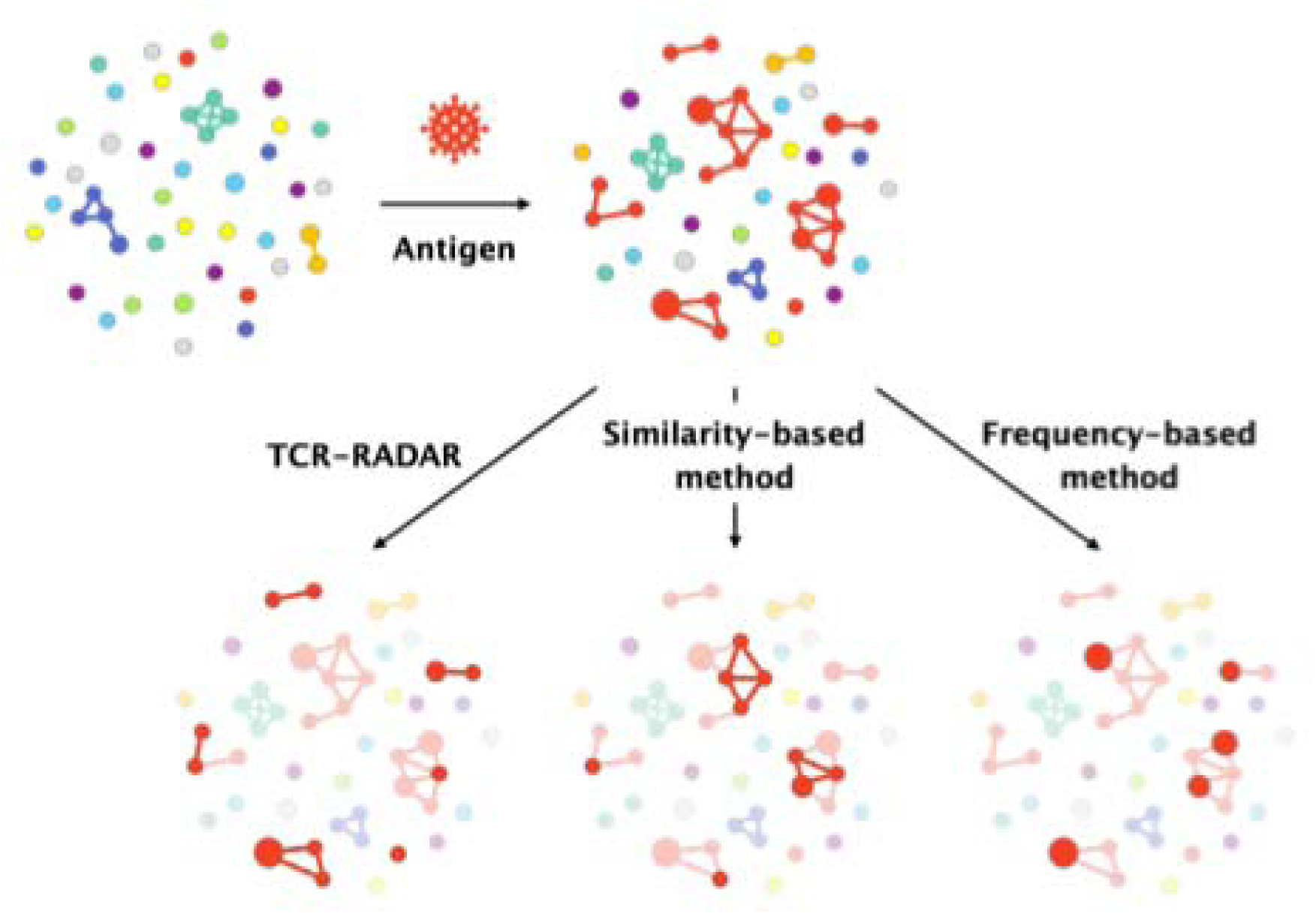
Complementary identification of antigen-specific clonotypes with proposed method, similarity-based method, and frequency-based method. Network visualization of TCR repertoire following antigen exposure, where nodes represent individual TCR clonotypes and edges connect sequence-similar TCRs. Node size reflects clonal frequency (clone count). Red nodes indicate antigen-specific TCRs. TCR-RADAR (left panel) uniquely identifies low-frequency antigen-specific TCRs at the periphery of sequence clusters, while similarity-based methods (middle panel) detect TCRs with enriched sequence neighbors, and frequency-based methods (right panel) capture highly expanded clones. The minimal overlap between methods demonstrates their complementary nature in comprehensive immune repertoire analysis. (The figure is adapted and redrawn from Ref. [21])

We validated TCR-RADAR on diverse immunological datasets including COVID-19 infection, influenza vaccination, and yellow fever (YF) vaccination. Our results demonstrate that this approach effectively identifies low-frequency antigen-specific TCRs that conventional methods fail to detect. TCR-RADAR establishes a new paradigm for antigen-specific TCR detection based on spatial distribution characteristics rather than sequence similarity or clonal frequency alone.

## Methods

### TCR-RADAR algorithm

TCR-RADAR identifies candidate antigen-specific TCRs by computing anomaly scores based on TCR sequence distances between two states: a’reference state’ (baseline condition) and a’query state’ (post-antigen exposure), identifying TCRs in the query state that are distant from the reference repertoire. Unlike frequency-based methods requiring multiple biological replicates, our approach operates with a single reference state and a single query state, reducing experimental complexity and enabling application to studies with limited sample availability. The algorithm consists of three main steps: preprocessing, anomaly score calculation, and candidate selection **(Fig. 2)**.

**Figure 2.**
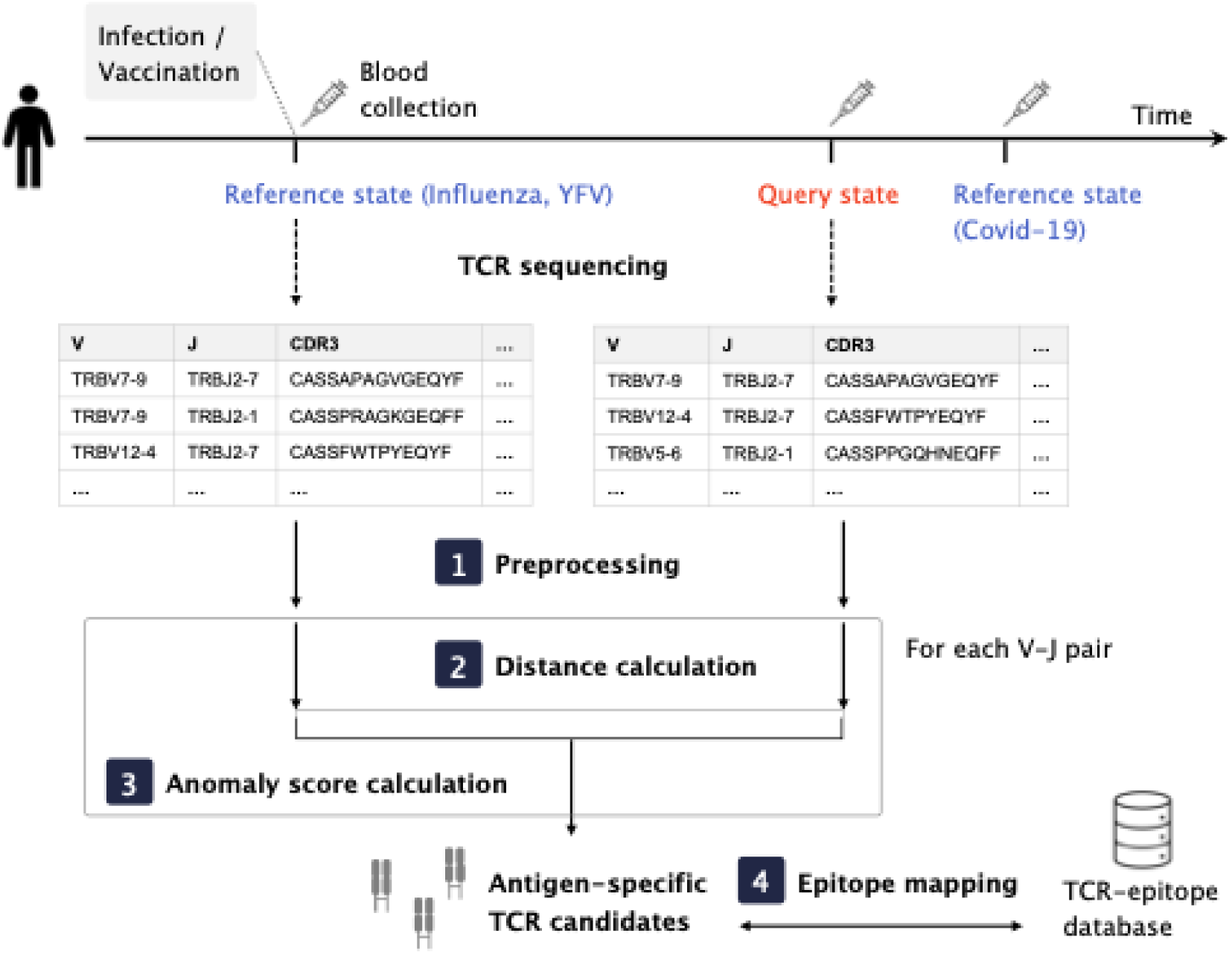
Schematic workflow of the distance-based anomaly detection method for identifying antigen-specific TCRs. TCR repertoires are sequenced from reference and query states following infection or vaccination. After preprocessing, TCRdist3 calculates pairwise distances within V-J gene combinations. Anomaly scores quantify the dissimilarity of query TCRs to the reference repertoire, with top-scoring candidates validated against TCR-epitope databases.

### Preprocessing of TCR repertoire data

TCR repertoire data from the reference and query states are processed as follows: (1) Non-functional V genes are removed using the functional V gene list from the tidytcells library [27]; (2) Non-productive sequences containing asterisks or underscores in the CDR3β regions are excluded; (3) Identical V-J-CDR3β sequences are collapsed by summing their clone counts; (4) TCRs with clone counts below a threshold (ranging from 1 to 10) are filtered out.

### Anomaly score calculation

To utilize potential distribution patterns of antigen-specific TCRs within TCR sequence space and to enable computationally efficient analysis of large repertoires, TCR-RADAR calculates anomaly scores within V or V-J gene groups. The granularity of gene-based grouping is adaptively determined by dataset size: V-J gene pairs are used when the total TCR count (combined from both reference and query datasets) exceeds 200,000; otherwise, V genes alone are used to maintain adequate sample sizes within each group.

Pairwise distances between TCR sequences are calculated using TCRdist3, generating two distance matrices: a reference-query matrix and a query-query matrix [18]. For each query TCR, the anomaly score is calculated as follows.

Let *Q_g_* and *R_g_* denote the sets of query and reference TCRs within a V-J gene combination *g*, respectively. For a query TCR *q_i_ ɛ Q_g_*, the base score is defined as the sum of TCRdist3 distances to all reference TCRs:

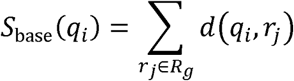

where d(q_i_,r_j_) represents the TCRdist3 distance between query TCR *q_i_* and reference TCR *r_j_*.

To incorporate local sequence context, the final anomaly score aggregates base scores from neighboring query TCRs:

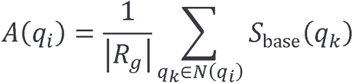

where the neighborhood *N*(*q_i_*) is defined as *N*(*q_i_*) ={*q_k_ ɛ Q_g_:d(q_i_,q_k_)<τ*}.

The radius threshold τ = 12.5 TCRdist units corresponds to a maximum distance of one amino acid mismatch in the CDR3 region. Normalization by |R_g_| ensures comparable scores across different V-J gene combinations with varying reference repertoire sizes.

### Candidate selection

Query TCRs were ranked by the anomaly score in descending order, with higher scores indicating greater dissimilarity from the reference repertoire, suggesting an increased likelihood of antigen specificity. The top 1,000 TCRs were selected as candidate antigen-specific sequences.

### Datasets

We evaluated our method using three TCR repertoire datasets from different immunological contexts, each representing distinct patterns of T cell clonal dynamics.

### COVID-19 infection dataset

We used publicly available TCRβ repertoire data from PBMCs of a male donor with mild COVID-19 [28]. T cell clonal frequencies following infection demonstrated three distinct patterns [28]. We focused specifically on the contracting clone population that expanded on day 15 post-infection and subsequently contracted by day 85. To capture these contracting clones, we selected day 85 post-infection as the reference state and day 15 post-infection as the query state.

### Influenza vaccination dataset

We analyzed TCRβ repertoire data from PBMCs of a donor who received influenza vaccination [26]. Day 0 (the day of vaccination) served as the reference state. We selected day 45 post-vaccination as the query state, representing when TCR clones that had significantly increased in frequency from day-14 reached their peak following vaccination.

### Yellow fever vaccination dataset

We utilized TCRβ repertoires from PBMCs of a S1 donor who received yellow fever (YF) vaccination (YFV17D) [25]. Day 0 (the day of vaccination) served as reference and day 15 post-vaccination as the query state, corresponding to peak TCR clonal expansion observed consistently across all six donors in the study.

### Validation of candidate TCRs using TCR-epitope database

To validate the antigen specificity of the unsupervisedly identified candidate TCRs, we utilized established TCR-epitope binding databases. For COVID-19 analysis, we obtained 154,320 CD8^+^ T cell-derived TCR-peptide binding pairs from the ImmuneCODE database [29], representing the largest validation dataset of the three disease contexts examined. For influenza analysis, we retrieved 6,669 TCR-peptide pairs from VDJdb [30] (accessed May 28, 2025). For the YF vaccination analysis in the main text, we validated using 1,851 NS4B-specific CD8^+^ T cells [31], while in the supplementary materials, we used 405 CD8^+^ T cells specific to NS4B_214-222_ LLWNGPMAV epitope from VDJdb (accessed May 26, 2025).

TCR sequences were considered matches based on the following criteria: For COVID-19, TCRs with identical V and J genes and at most one amino acid difference in the CDR3 region were classified as highly similar. TCRs were considered epitope-specific when at least two highly similar sequences recognized the same epitope. For influenza and YF datasets, matching required identical V and J genes and allowed up to two amino acid mismatches in CDR3 sequences.

The validation accuracy was defined as the proportion of candidate TCRs successfully mapped to known epitope-specific sequences in the respective databases.

### Identification of antigen-specific clones using frequency-based and similarity-based methods

To benchmark our approach, we compared its performance against established frequency-based and similarity-based methods for identifying antigen-specific TCRs.

### Frequency-based methods

We employed two frequency-based approaches that detect antigen-specific TCRs through statistical analysis of clonal expansion between reference and query states. First, we applied edgeR [24], originally developed for differential gene expression analysis in RNA-seq experiments and successfully adapted for detecting TCR clonal expansion [25,26,28,31,32]. The edgeR method requires two biological replicates for both reference and query states.

Biological replicates at each time point were combined, and clones with average counts below 4 were filtered to improve statistical power. Library sizes were normalized using the Trimmed Mean of M-values (TMM) method, and dispersion parameters were estimated using quantile-adjusted conditional maximum likelihood (qCML). Significant expansion was defined as log2 fold change ≥5 with FDR ≤0.05 for COVID-19 and YF datasets, and log2 fold change ≥3 with FDR ≤0.05 for influenza data.

Second, we employed Pogorelyy’s method, a Bayesian statistical framework specifically designed for detecting TCR clonal expansion and contraction [25]. This method requires two biological replicates for the reference state and one for the query state. We applied the same fold change thresholds as edgeR with P-value ≤0.05 as the significance threshold.

### Similarity-based method

We also evaluated ALICE, which identifies antigen-specific TCRs from single snapshot data without requiring longitudinal samples [21]. ALICE employs a statistical model of V(D)J recombination to generate synthetic amino acid sequences, then detects TCRs with significantly enriched neighbors within CDR3 sequence space compared to expected values. We configured ALICE with the following parameters: cores=20, iter=10, and nrec=5×10^5^, yielding 100 million total simulated sequences. Computations were performed on an AMD Ryzen Threadripper PRO 5995WX system (64 cores/128 threads, 256 MB cache) with 512 GB ECC Registered DDR4-3200 memory.

All candidate TCRs identified by these methods were validated against epitope databases following the same mapping procedure as our proposed method to ensure fair accuracy comparison.

### Use of artificial intelligence tools

Large language models were used during manuscript preparation as an aid for correcting written text, for assisting in code writing, and for creating code documentation. All AI-generated outputs were reviewed, verified, and edited by the authors, who take full responsibility for the content of this work.

## Results

### TCR-RADAR captures antigen-specific TCRs at V gene cluster peripheries

To characterize the spatial distribution of antigen-specific TCRs in sequence space, we first visualized the TCR repertoire from a donor infected with COVID-19 [28]. t-SNE visualization revealed distinct clusters corresponding to different V gene families **(Fig. 3a)**. When we mapped SARS-CoV-2 epitope-specific TCRs onto this visualization, they predominantly localized to the peripheries of V gene clusters rather than cluster centers **(Fig. 3b)**. This peripheral distribution provided the rationale for our anomaly detection approach: antigen-specific TCRs occupy unique positions in sequence space within their respective germline gene groupings, making them well-suited for distance-based anomaly detection.

**Figure 3.**
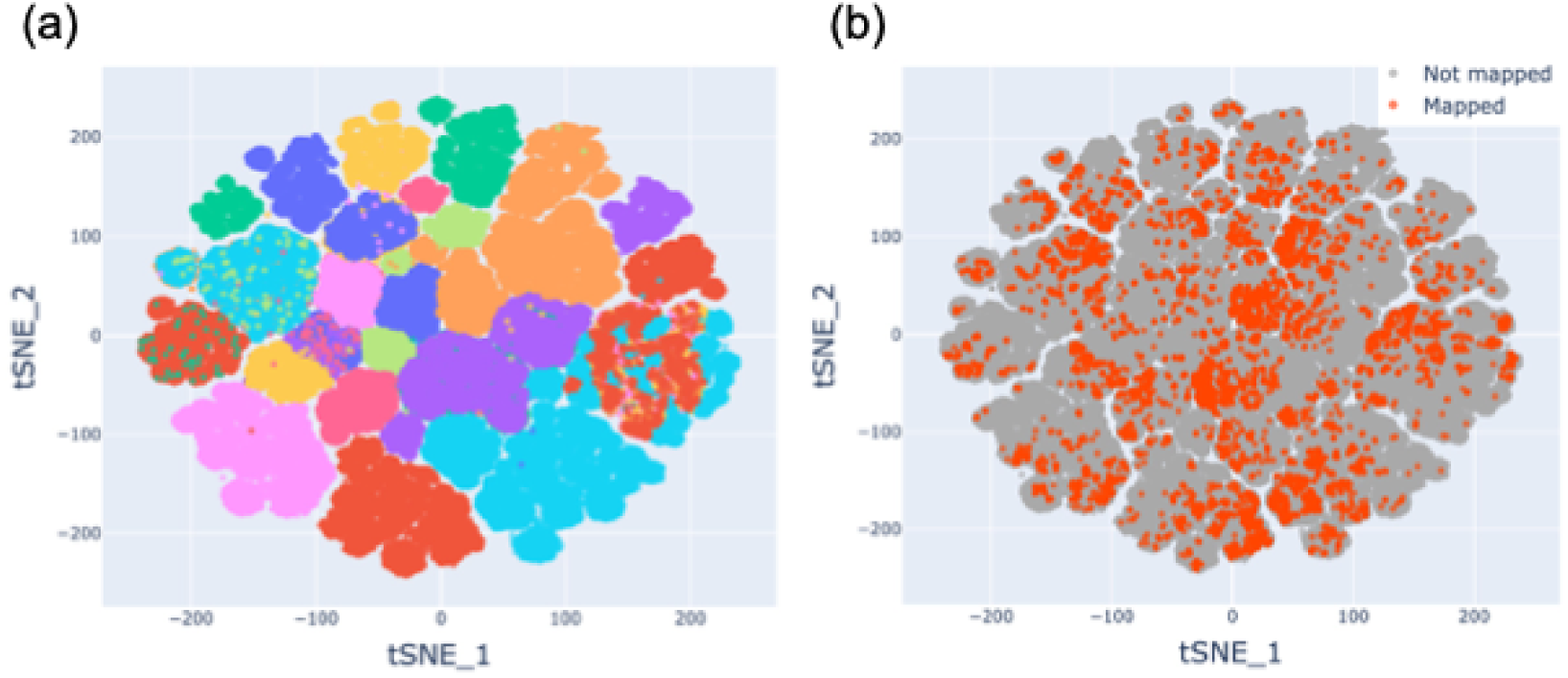
Spatial distribution of TCR repertoires reveals peripheral localization of SARS-CoV-2-specific TCRs. (a) t-SNE visualization of TCR repertoire from COVID-19 infected donor at day 15 post-infection (query state). Latent representations were obtained using the protein language model SCEPTR [45] and projected onto two dimensions via t-SNE; distinct clusters correspond to different V gene families. (b) Same visualization with SARS-CoV-2 epitope-specific TCRs highlighted in red, demonstrating preferential localization at V gene cluster peripheries rather than cluster centers, supporting the rationale for distance-based anomaly detection.

### Superior performance in detecting low-frequency SARS-CoV-2-specific TCRs

We first evaluated our method on COVID-19 infection data, validated against a comprehensive dataset of over 150,000 experimentally confirmed SARS-CoV-2-specific TCRs. TCR-RADAR achieved 34.3% accuracy in identifying antigen-specific TCRs, significantly outperforming both similarity-based (ALICE: 8.0%) and frequency-based methods (edgeR: 5.8%, Pogorelyy: 6.3%) **(Table 1)**. Given the large dataset (>200,000 TCRs), calculating pairwise distances for all pairs is computationally challenging. TCR-RADAR circumvents this problem by applying V-J gene pair grouping when calculating anomaly scores, which enables efficient processing in 23 minutes on a computer only with 16GB RAM. Notably, the overlap between TCRs identified by our method and those identified by existing approaches was minimal (0.3-0.6%), indicating that TCR-RADAR captures a distinct population of functionally important TCRs overlooked by conventional analyses.

**Table 1.** Comparative analysis of antigen-specific TCR detection methods using COVID-19 infection data. TCR-RADAR achieved 34.3% accuracy with minimal overlap with similarity-based (ALICE) and frequency-based (edgeR, Pogorelyy) approaches, detecting TCRs at clone count 1. Row headers indicate: Required sample (total number of samples needed at reference and query states with replicate numbers in parentheses), Computational time (runtime in minutes), Antigen-specific TCRs (mapped TCRs/candidate TCRs with accuracy percentage), Overlap (percentage of mapped TCRs overlapping with TCR-RADAR), and Minimum clone count (lowest detection threshold for mapped TCRs).

|  | TCR-RADAR | ALICE | edgeR | Pogorelyy |
| --- | --- | --- | --- | --- |
| Required sample | 2 (Day85: 1 replicate, Day15: 1 replicate) | 1 (Day15: 1 replicate) | 4 (Day85: 2 replicate, Day15: 2 replicate) | 3 (Day85: 2 replicate, Day15: 1 replicate) |
| Computational time (min) | 23 | 300 | < 1 | 111 |
| Antigen-specific TCRs | 343/1000 (34.3%) | 47/586 (8.0%) | 65/1113 (5.8%) | 34/536 (6.3%) |
| Overlap (%) | - | 0.6 | 0.6 | 0.3 |
| Minimum clone count | 1 | 1 | 8 | 20 |

A key advantage of our approach is its ability to detect low-frequency TCRs. While frequency-based methods could only identify TCRs with minimum clone counts of 8 (edgeR) and 20 (Pogorelyy method), our approach successfully detected antigen-specific TCRs with clone counts as low as one **(Table 1**, **Fig. 4a)**. The clone count distribution of mapped TCRs (candidate TCRs which are validated against experimentally confirmed databases) demonstrated that our method consistently identified lower-frequency clones compared to frequency-based approaches **(Fig. 4a)**.

**Figure 4.**
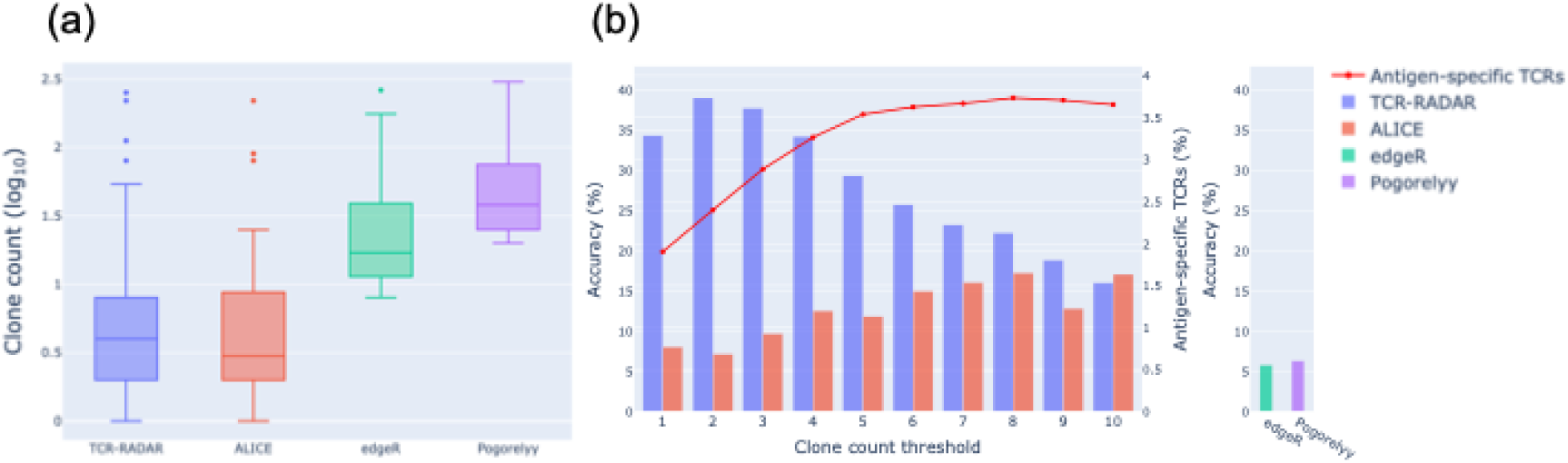
Low-frequency TCR detection performance for COVID-19 infection data. (a) Clone count distributions of mapped TCRs identified by each method. (b) Accuracy comparison across different clone count thresholds. TCR-RADAR and ALICE were evaluated at each threshold, while frequency-based methods (edgeR and the Pogorelyy method) were evaluated on the entire TCR repertoire. The red line indicates the fraction of antigen-specific TCRs at each threshold.

To understand the relationship between clone frequency and detection accuracy, we evaluated performance across different clone count thresholds **(Fig. 4b)**. TCR-RADAR maintained high accuracy (>30%) even at the lowest clone count thresholds, where the fraction of antigen-specific TCRs showed a stepwise decline below a threshold of 4. In contrast, existing methods generally showed lower accuracy across the tested thresholds, where frequency-based methods were evaluated only on the entire TCR repertoire. Interestingly, when the fraction of antigen-specific TCRs plateaued at clone counts of 5 or greater, TCR-RADAR’s accuracy showed a gradual decline, suggesting optimal performance in the low-frequency range where conventional approaches exhibit limited sensitivity.

### Robust detection independent of clonal expansion magnitude in influenza vaccination

To evaluate our method’s performance under conditions of limited immune response, we analyzed TCR repertoires from influenza vaccination, where clonal expansion is limited [26]. Sycheva et al. showed that changes in T cell clone frequency on days 5 and 12 post-vaccination were comparable to baseline variations between days-14 and 0, presenting a challenging scenario for frequency-based methods.

Despite these constraints, TCR-RADAR achieved 22.5% accuracy, significantly higher than ALICE (7.2%) and frequency-based approaches (edgeR: 5.3%, Pogorelyy: 0%) **(Table 2)**. Notably, the Pogorelyy method failed to identify any epitope-specific TCRs, demonstrating the limitations of frequency-based approaches when clonal expansion is minimal. The minimal overlap between TCR-RADAR and others (ALICE: 6.7%, edgeR: 0%) further indicated that TCR-RADAR captures distinct TCR populations **(Table 2)**.

**Table 2.**
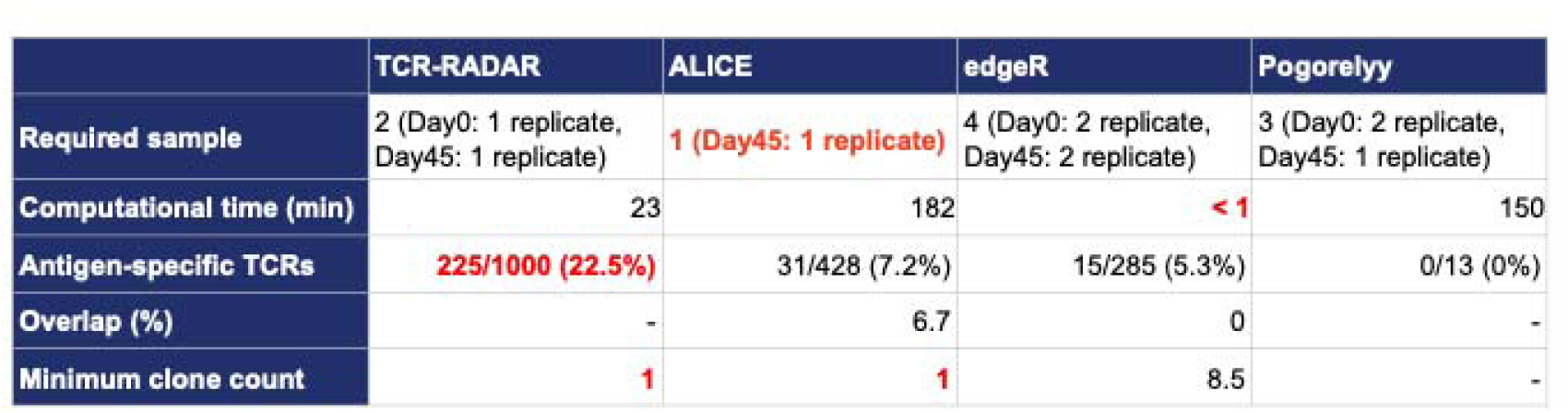
Performance comparison on influenza vaccination data with limited clonal expansion. TCR-RADAR maintained 22.5% accuracy while the Pogorelyy method failed to identify any epitope-specific TCRs, demonstrating robustness under limited clonal expansion conditions.

TCR-RADAR detected influenza-specific TCRs with clone counts as low as one, while edgeR required average minimum counts of 8.5 **(Fig. 5a)**. Across clone count thresholds 1-6, TCR-RADAR’s accuracy tracked closely with the fraction of antigen-specific TCR, maintaining consistently higher accuracy than ALICE at all thresholds **(Fig. 5b)**. These results demonstrate that our approach remains effective even when antigen-driven proliferation is limited.

**Figure 5.**
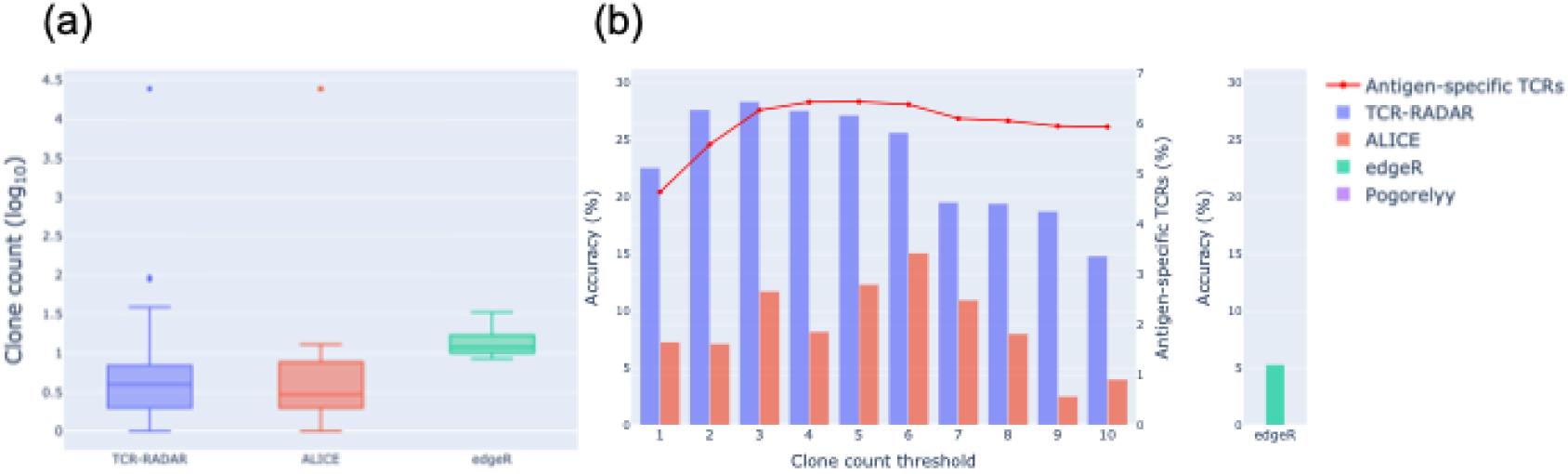
Low-frequency TCR detection performance for influenza vaccination data. (a) Clone count distributions of mapped TCRs identified by each method. (b) Accuracy comparison across different clone count thresholds. The red line indicates the fraction of antigen-specific TCRs at each threshold.

### Performance characteristics in strong immune responses to yellow fever vaccination

To evaluate our method under conditions of robust immune response, we analyzed TCR repertoires following YF vaccination, which reproducibly induces T cell responses across different donors, consistently peaking around 2 weeks post-vaccination, followed by contraction and long-term memory formation [25,31,33,34]. Although the overall accuracy of TCR-RADAR (15.6%) was lower than that of ALICE (29.4%) and frequency-based methods (edgeR: 17.1%, Pogorelyy: 18.5%) **(Table 3)**, the minimal overlap with other methods (ALICE: 1.3%, edgeR: 1.3%, Pogorelyy: 0.6%) indicates that TCR-RADAR detects complementary TCR populations.

**Table 3.** Performance evaluation on yellow fever vaccination data. ALICE achieved highest accuracy (29.4%) while TCR-RADAR showed complementary detection with minimal overlap (≤1.3%).

|  | TCR-RADAR | ALICE | edgeR | Pogorelyy |
| --- | --- | --- | --- | --- |
| Required sample | 2 (Day0: 1 replicate, Day15: 1 replicate) | 1 (Day15: 1 replicate) | 4 (Day0: 2 replicate, Day15: 2 replicate) | 3 (Day0: 2 replicate, Day15: 1 replicate) |
| Computational time (min) | 29 | 223 | < 1 | 29 |
| Antigen-specific TCRs | 156/1000 (15.6%) | 144/490 (29.4%) | 246/1436 (17.1%) | 141/762 (18.5%) |
| Overlap (%) | - | 1.3 | 1.3 | 0.6 |
| Minimum clone count | 1 | 1 | 8 | 17 |

Furthermore, TCR-RADAR maintained its core advantages in detecting low-frequency clones, identifying YF-specific TCRs with clone counts of 1, compared to minimum counts of 8 (edgeR) and 17 (Pogorelyy) for frequency-based methods **(Fig. 6a)**. Notably, within the clone count threshold ranging from 2 to 8, TCR-RADAR achieved comparable accuracy to frequency-based approaches while successfully detecting lower-frequency TCRs **(Fig. 6b)**. Validation using an independent dataset of 405 NS4B_214-222_-specific T cells from VDJdb confirmed the superior performance of ALICE **(Fig. 7**, **Table 4)**.

**Figure 6.**
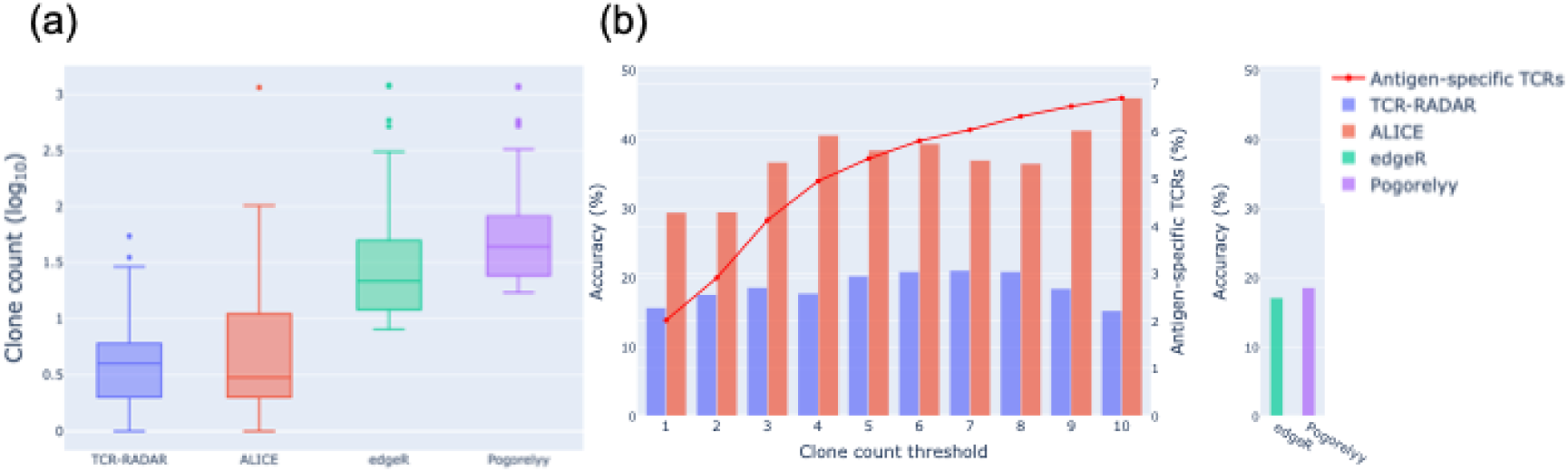
Low-frequency TCR detection performance for yellow fever vaccination data. (a) Clone count distributions of mapped TCRs identified by each method. (b) Accuracy comparison across different clone count thresholds. The red line indicates the fraction of antigen-specific TCRs at each threshold.

**Figure 7.**
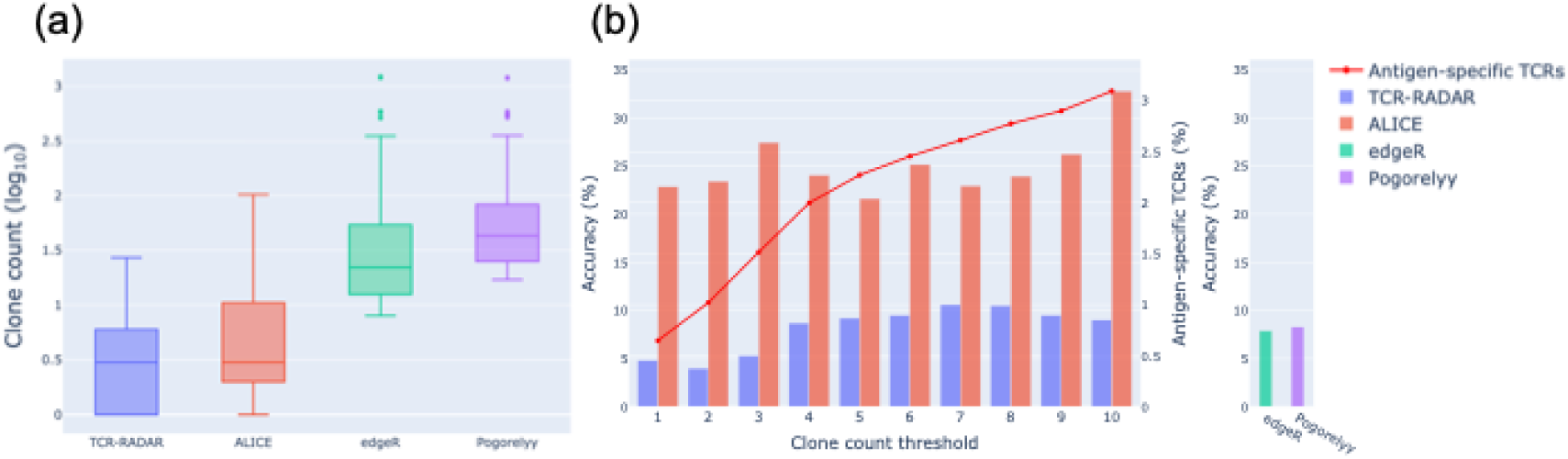
Low-frequency TCR detection performance for yellow fever vaccination data validated against VDJdb dataset. (a) Clone count distributions of mapped TCRs identified by each method. (b) Accuracy comparison across different clone count thresholds. The red line indicates the fraction of antigen-specific TCRs at each threshold.

**Table 4.** Performance evaluation on yellow fever vaccination data validated against VDJdb dataset. Validation against VDJdb NS4B_214-222_ epitope-specific TCRs (n=405) revealed consistent patterns observed in the main analysis with Minervina dataset (Table 3): ALICE achieves highest accuracy (22.9%), while TCR-RADAR (4.8%) maintains its unique ability to detect TCRs at clone count 1 with minimal overlap (≤6.3%) with conventional approaches.

|  | TCR-RADAR | ALICE | edgeR | Pogorelyy |
| --- | --- | --- | --- | --- |
| Required sample | 2 (Day0: 1 replicate, Day15: 1 replicate) | 1 (Day15: 1 replicate) | 4 (Day0: 2 replicate, Day15: 2 replicate) | 3 (Day0: 2 replicate, Day15: 1 replicate) |
| Computational time (min) | 49 | 223 | < 1 | 29 |
| Antigen-specific TCRs | 48/1000 (4.8%) | 112/490 (22.9%) | 113/1436 (7.9%) | 63/762 (8.3%) |
| Overlap (%) | - | 6.3 | 2.1 | 0 |
| Minimum clone count | 1 | 1 | 8 | 17 |

## Discussion

We developed TCR-RADAR, an unsupervised approach for detecting low-frequency antigen-specific TCRs through distance-based anomaly detection, distinct from conventional frequency-based and similarity-based methods. TCR-RADAR employs TCRdist3 to identify TCRs that are anomalous compared to reference repertoires within the clusters of V-J gene pairs or V gene alone, based on our observation that antigen-specific TCRs localize at the peripheries of V gene clusters in sequence space **(Fig. 3)**. TCR-RADAR’s strategy of computing distances within each gene cluster significantly reduces computational costs, enabling the analysis of massive TCR repertoire datasets exceeding 1 million sequences on computing resources with 16GB RAM, making large-scale immunological analyses practically feasible. This approach extends the current methodological framework for TCR specificity detection beyond the traditional dichotomy of frequency-based versus similarity-based methods, introducing a novel “anomaly-based” paradigm. The spatial distribution of antigen-specific TCRs at cluster peripheries provides new insights into TCR repertoire organization that could inform future algorithm development for TCR specificity prediction.

TCR-RADAR specifically targets low-frequency antigen-specific TCRs, which are difficult to detect using existing methods due to their algorithmic limitations. The frequency distribution of TCRs follows a power law [7,8], with antigen-specific TCRs occurring at extremely low frequencies of approximately one per million sequences [9–11]. In our analysis, TCRs with a clone count of 1 represented a substantial proportion of antigen-specific populations (COVID-19: 44.9%, Influenza: 66.1%, YF: 68.6%), highlighting the importance of detecting these rare clones. Furthermore, technical factors such as PCR amplification bias and sequencing errors can artificially reduce the apparent frequency of truly abundant clones [35–37]. Therefore, frequency-independent detection methods are essential for accurate and comprehensive analysis of antigen-specific TCR repertoires.

TCR-RADAR complements experimental validation methods. The most common experimental approach for identifying antigen-specific T cells, the MHC multimer assay [38], requires prior knowledge of MHC restriction and epitope sequences. Due to the extremely low frequency of antigen-specific T cells, these assays necessitate either magnetic bead enrichment techniques or analysis of large sample volumes, imposing significant costs [39]. Consequently, existing antigen-specific TCR databases are biased toward well-characterized epitopes [40,41]. Given the scarcity of known TCR-antigen training pairs, computational models have not yet achieved complete mapping of unseen epitopes [15]. TCR-RADAR prioritizes TCR candidates for experimental validation, potentially reducing experimental costs and effort while accelerating the expansion of antigen-specific TCR databases needed for robust machine learning model evaluation.

The lower performance on YF data warrants further investigation. It is possible that YF-induced immune responses differ fundamentally from other infections. Unlike COVID-19 and influenza datasets, ALICE achieved the highest accuracy in detecting YF-specific TCRs, identifying TCRs with enriched sequence neighbors compared to V(D)J recombination predictions. Previous studies have demonstrated limited diversity in YF-specific TCR responses, particularly a strong bias toward specific TCRα chains [31,42]. This constrained diversity suggests that YF vaccination induces highly convergent immune responses across individuals, enabling generative models like ALICE to effectively capture YF-specific TCRs through sequence similarity, while COVID-19 and influenza may elicit more diverse, individualized responses. Future development of integrated detection systems that combine complementary properties of TCR-RADAR with similarity-based and frequency-based approaches could elucidate the principles governing antigen specificity. Alternatively, limitations in the validation databases may explain the performance difference. Our evaluation relies on known TCR-epitope pairs from public databases (ImmuneCODE, VDJdb, and published datasets), potentially misclassifying true antigen-specific TCRs absent from these databases as false negatives. The YF validation set contains only 1,851 TCR pairs compared with 154,320 pairs in COVID-19 and 6,669 in influenza vaccination, and this small and potentially biased dataset may not accurately reflect true performance of validated methods. Expansion of YF epitope-specific TCR datasets would help resolve this uncertainty.

Our study has several limitations. Although TCR-RADAR is an unsupervised approach, it relies on three hyperparameters: the TCR count threshold (200,000) for gene-based grouping, the neighboring radius (12.5 TCRdist units), and the number of selected candidates (1,000). While tuning these values for each dataset could enhance performance, the limited availability of validation data led us to adopt fixed defaults—the radius was chosen to reflect the distance of a single amino acid substitution and the candidate number was chosen to be comparable to existing methods. Additionally, while our COVID-19 validation employed stringent criteria requiring at least two similar TCRs (matching V-J genes with ≤1 CDR3 mismatch) per epitope due to the large database size, the smaller influenza and YF databases required more relaxed mapping criteria (matching V-J genes with ≤2 CDR3 mismatches).

This relaxation is supported by evidence that similar sequences often share antigen specificity [16,17,43], but experimental validation remains essential to confirm binding. Furthermore, our analysis focused solely on TCRβ chains from bulk repertoire data. Since pMHC recognition involves both α and β chains, incorporating TCRα information should improve prediction accuracy [44]. As TCRdist3 supports paired-chain data, our method can be readily extended to single-cell datasets. However, validation using longitudinal single-cell data following infection or vaccination is currently limited by data availability. These constraints—ranging from hyperparameter optimization to validation criteria to paired-chain analysis—all stem from the fundamental scarcity of comprehensive TCR-epitope datasets.

Future expansion of TCR-epitope binding pair databases will facilitate both systematic hyperparameter optimization and more robust validation.

## Conclusions

We present a fundamentally new paradigm for detecting antigen-specific TCRs through distance-based anomaly detection, moving beyond the traditional dichotomy of frequency-based and similarity-based approaches. Our key observation that antigen-specific TCRs are preferentially localized at the periphery of V gene clusters in TCR sequence space led to the development of an unsupervised method that successfully identifies low-frequency TCRs missed by conventional approaches. Across three diverse immunological contexts—COVID-19 infection, influenza vaccination, and yellow fever vaccination—TCR-RADAR demonstrated its unique capability to detect antigen-specific TCRs at clone count one, achieving 34.3% accuracy for SARS-CoV-2-specific TCRs, whereas existing methods were limited to detecting higher-frequency clones. The minimal overlap (≤6.7%) with conventional approaches confirms that TCR-RADAR captures distinct TCR populations, providing a complementary tool for comprehensive immune repertoire analysis. Our findings suggest that antigen-specific TCR distributions vary by pathogen type—some converging toward similar sequences while others localizing as anomalies—offering new insights into the fundamental diversity of immune recognition patterns. As TCR sequencing technologies advance and epitope databases expand, TCR-RADAR provides a valuable strategy for prioritizing rare antigen-specific clones for experimental validation, potentially accelerating the development of diagnostic tools and immunotherapies.

## Acknowledgements

We are grateful to Mikhail V. Pogorelyy and Anastasiia L. Sycheva for sharing processed influenza and YFV data and for their helpful comments on our analysis of the validation datasets. The first author is supported by the World-leading Innovative Graduate Study Program Co-designing Future Society (WINGS-CFS), The University of Tokyo.

## Data and Code Availability

All data analyzed in this study are publicly available. The TCRβ repertoire datasets were derived from sources in the public domain: Minervina et al. [28] for COVID-19 infection, Sycheva et al. [26] for influenza vaccination, and Pogorelyy et al. [25] for yellow fever vaccination. TCR-epitope binding data were obtained from the ImmuneCODE database (https://clients.adaptivebiotech.com/pub/covid-2020) [29] and VDJdb (https://vdjdb.cdr3.net) [30].

